# Bidirectional Cycling Dynamics of Living Neuronal Networks *in Vitro*

**DOI:** 10.1101/223230

**Authors:** Arseniy Gladkov, Oleg Grinchuk, Pigareva Yana, Irina Mukhina, Victor Kazantsev, Alexey Pimashkin

## Abstract

Phenomena of synchronization, rhythmogenesis and coherence found in brain networks are believed to be a dynamical substrate for cognitive functions such as learning and memory. However, it is still debated whether the rhythmic activity emerges from network morphology developed in neurogenesis or as a result of neuronal dynamics realized under certain conditions. In this research we found, that in neural networks formed in mature hippocampal cultures with high cellular density the spiking activity self-organized and converged to long, complex and rhythmically repeated superbursts. The superburst lasted tens of seconds and consisted of hundreds of short (50-100 ms) small bursts with a high spiking rate of 139.0 ± 78.6 Hz that can be associated with high-frequency oscillations in the hippocampus. In turn, the interval between peak burst activities in the range of 100-150 ms can be treated as a theta rhythm (11.2 ± 1.5 Hz). Distribution of spikes within the bursts was non-random, representing a set of well-defined space-time base patterns or motifs. We found that the long superburst can be classified into two types. Each type was associated with a unique direction of spike propagation and, hence, can be encoded by a binary sequence with random switching between the two “functional” states. Such precisely structured bidirectional rhythmic activity developed in self-organizing cultured networks were quite similar to what observed in the *in vivo* experiments.

## 1. Introduction

Synchronization and interplay between excitation and inhibition in neural networks play crucial role in brain rhythmic activity organization [1–5]. Rhythmic oscillatory activity with various frequencies represents a multi-clock substrate for cognitive function, memory and sleep [6, 7]. It is, however, still a question whether the rhythmicity emerged from specific network morphology developed in neurogenesis [7–11] or it can be generated spontaneously due to network nonlinear dynamics mediated by an interplay of excitation-inhibition and sustained by homeostatic balance [12–14]. An answer to this fundamental question promises to define network mechanisms of pathological seizure activity and, hence, to determine treatment approaches. Recent studies have shown that many brain network functions, normal and pathological states can be studied using *in vitro* models [13–23]. In this aspect dissociated neuronal cultures gives us unique possibility to model network dynamics and rhythmicity *in vitro*.

It has been shown in several studies that dissociated neuronal cultures plated on microelectrode arrays after several days *in vitro* (DIV) spontaneously generated activity in the form of periodic and synchronized network burst discharges [24-26,15]. The bursts had various space-time distributions of spikes recorded at the electrodes during the discharges. It has been discussed that network bursts may be involved in mechanisms of information encoding [16], memory [17] and chronic neurological diseases, such as epilepsy [18, 27]. For example, similar burst dynamics is developed spontaneously or evoked by stimulus *in vivo* in the cortex, hippocampus, and brain nuclei during brain development [19, 20, 28, 29]. Such *in vivo* bursts were associated with a single sharp potential or spindle-shaped field oscillations (approximately 10 Hz) [19].

Regular bursts in cultured networks were characterized by variable firing rates. At the same time, they were composed of highly precise and reproducible spatio-temporal spiking patterns. The spiking patterns can be quantified by the values of timings and the recruitment order of the first spikes initiating the bursts for each electrode, called activation patterns [30]. Such patterns were reported to be stable on a timescale of several hours [30–32]. It has also been shown that the profile of the spiking patterns (several tens of milliseconds) at the beginning of the bursts was precisely repeated in the subsequent bursts, while the middle phase of burst formation was highly variable [31]. Analysis of spontaneous activity in the cultured networks grown on high-density microelectrode arrays (4096 electrodes) also revealed that only short intervals in the initial parts of the bursts were reproducible and could be associated with spike propagation in the network from certain initiation points (neurons) [33]. Detailed analysis showed that bursting activity consisted of several motifs that can be distinguished by direction or spike propagation pathways and that appear randomly during the recording. Several types of patterns, i.e., motifs, were also found in the spontaneous bursting activity using activation patterns (only first spike timings) and spiking frequency patterns [21, 34].

The bursting activity in neuronal cultures changed dramatically during development *in vitro* and essentially depended on initial cell plating density [26, 35]. The minimum plating density of a cortical culture required for bursting activity to emerge was found to be 250 cells per mm^2^ in Neurobasal medium (neuron culture medium) [36]. During the first 3 weeks of development, the numbers of GABAergic and glutamatergic terminals increased gradually, simultaneously with the bursting rate [36]. A steady state reached after 3-4 weeks in vitro in hippocampal [37–39] and in cortical [36, 40, 41] cultures. In the mature stages of highly dense cultures, the spiking activity consisted of complex sequences of a type of burst often called superburst, showing durations from several to tens of seconds. The superburst were observed only in dense dissociated cultures (2500 rat cortical cells per mm^2^ in Dulbecco’s modified Eagle’s medium (DMEM) [22, 26], 4000 cells per mm^2^ in Neurobasal medium only with blockers of inhibitory connections [18] or 8x10^3^ and 10^6^ rat hippocampal cells per mm^2^ in DMEM [42, 43]). The overall network activity was very complex and characterized by spontaneous superbursts which, in turn, may cluster into small superburst series [44]. The superbursting activity was also found in multilayered neural cultures [45]. High-frequency oscillations resembling network superbursts were observed even in small but dense neuronal clusters (up to 40 cells) [35]. Numerical simulation of cultured networks revealed that superbursts in larger networks (up to 50 000 neurons) occurred at earlier stages of network development comparing to smaller networks (up to 11 000 neurons) [46]. So that, the network size could be also a crucial parameter [46].

Superbursts in neuronal cultures represented highly coherent spatio-temporal spiking activity patterns spontaneously developed in originally nonstructured networks due to self-organization and plasticity [22].

One of the mechanisms suggested for superburst was the interaction between the activity of excitatory and inhibitory neurons in culture. In particular, the addition of inhibitory cells from the striatum to the hippocampal culture was used to study their impact to burst dynamics. Increase of inhibitory cell fraction from 20% to 56% in neuronalculture significantly increased the number of small bursts in the superburst structure [47]. The important role of GABAergic neurons in bursting activity generation was also confirmed in network mathematical models which GABAergic neurons were involved in the small burst generation in subsequent superburst [46]. Superbursting activity has been increased by blockers of inhibitory synapses (bicuculline, picrotoxin) or a low concentration of Mg^2+^ ions [18, 48] and has been decreased by a combination of Na^+^ channel blockers with picrotoxin [18].

The investigation of spiking patterns in the superbursts revealed remarkably precise repetition of the internal bursting sequence [22]. Superbursts appeared with irregular intervals, but their internal structure contained small bursts with highly regular and reproducible activation patterns during hours or days [26]. In another study of cortical cultures during the mature stage (4-6 week *in vitro)*, definite motifs in the burst activation pattern corresponded to specific oscillation phase during the ultra-slow oscillations (<0.01 Hz) was found [12]. Such ultra-slow oscillations could also be treated as superbursts with regular intervals. Spiking pattern motifs were found to be strongly conserved across multiple oscillation cycles, repeating themselves with high spatio-temporal precision.

One recent mathematical model predicted that high neurite and synaptic density may also influence on small bursts in the superburst subsequences [46]. Other studies showed that stable rhythmic activity in the form of propagating synchronized bursts during several minutes can be induced in cortical cultures under inhibition of GABAergic synaptic transmission [13]. Such periodic synchronized activity at 3-4 seconds time scale was observed only at the boundaries of the culture. Thus, the balance of excitation-inhibition should be an important parameter to generate in networks stable and reproducible synchronized activity.

In this study, we observed long superbursting activity with well-defined and reproducible temporal dynamics in spontaneously developed hippocampal culture on the microelectrode array (MEA). Regarding electrophysiological activity, we found long (up to 30 seconds) superburst consisting of subsequences of (up to hundreds) highly reproducible short bursts in the center of the cultured network. The spiking frequency in the bursts was found to be 139.0 ± 78.6 Hz, and interburst interval was in the range of 100-150 ms (11.2 ± 1.5 Hz), which resembled unique hippocampal activity in *in-vivo* conditions [49]. Spike propagation pathways during short bursts in all long superburst were aligned along two major spatial directions. We found that the long superburst could be encoded into two types, each associated with a definite orientation of spike propagation. The orientation was switched during each single superburst; in the following superburst, the orientation was determined at random, with a probability of switching to the next orientation at approximately 50%. Therefore, the superburst time sequence can be encoded by binary symbols reflecting spontaneous activation of two dominant spike propagation patterns selected in mature networks of cultured hippocampal cells. Note that such well-organized rhythmic activity emerged spontaneously in matured culture networks *in vitro* without any specific stimulation and afferentation. We believe that such self-organizing dynamics in culture development led to a certain balance of excitation and inhibition, where cycling dynamics serves as a homeostatic functional state of the culture. It also gives possible mechanism how different functional states (like, vortices and synchronized epileptic-like discharges) may spontaneously appear in brain network without any stimulation.

## 2. Materials and Methods

### 2.1 Cell culture

Hippocampal cells were dissociated from embryonic mice (E18) and plated on microelectrode arrays (MEAs) pre-treated with adhesion promoting molecules of polyethyleneimine (Sigma P3143) with a final density of approximately 15,000–20,000 cells/mm^2^ (Fig 1). Our cultures were composed of 4-5 layers of the cells. The mice used in our study were received from Institute of Bioorganic Chemistry *Pushchino*, Moscow Region, Russia. C57B1/6 mice were euthanized via cervical dislocation according to protocols approved by the National Ministry of Public Health for the care and use of laboratory animals. The protocol was approved by the Committee on the Ethics of Animal Experiments of the Nizhny Novgorod State Medical Academy (Permit Number: 9 - 25.09.2014). All efforts were made to minimize suffering. Embryos were removed and decapitated. The entire hippocampus was dissected under sterile conditions. The cortex, whole medulla and the lower part of the pons were excluded during the dissection. Hippocampi were cut in Ca^2+^- and Mg^2+^-free phosphate-buffered saline (PBS-minus). After enzymatic digestion for 25 min using 0.25% trypsin (Invitrogen 25200-056) at 37°C, cells were separated by trituration (10 passes) using a 1 ml pipette tip. Next, the solution was centrifuged at 1500 g for 1.5 min, and the cell pellet was immediately resuspended in Neurobasal culture medium (Invitrogen 21103-049) with B27 (Invitrogen 17504-044), glutamine (Invitrogen 25030-024) and 10% fetal calf serum (PanEco K055). The dissociated cells were seeded in a 30 μl droplet covering the centre of the culture dish with 1 mm^2^ electrode region of the MEA. It resulted to culture size of 6-7 mm in diameter. After the cells had adhered (usually in 2 hrs), the dishes were filled with 1 ml Neurobasal medium (NBM) supplemented with B-27 and 0.5 mM glutamine with 10% fetal calf serum. After 24 hrs, the plating medium was replaced with a medium containing NBM 2% B-27 and 1% glutamine and 0.5% fetal calf serum but with no antibiotics or antimycotics. Glial growth was not suppressed, given that glial cells are essential for long-term culture health. Half of the medium was replaced every 2 days. The cells were cultured under constant conditions of 35.5°C, 5% CO2 and 95% air at saturating humidity in a cell culture incubator (MCO-18AIC, SANYO).

**Fig 1.**
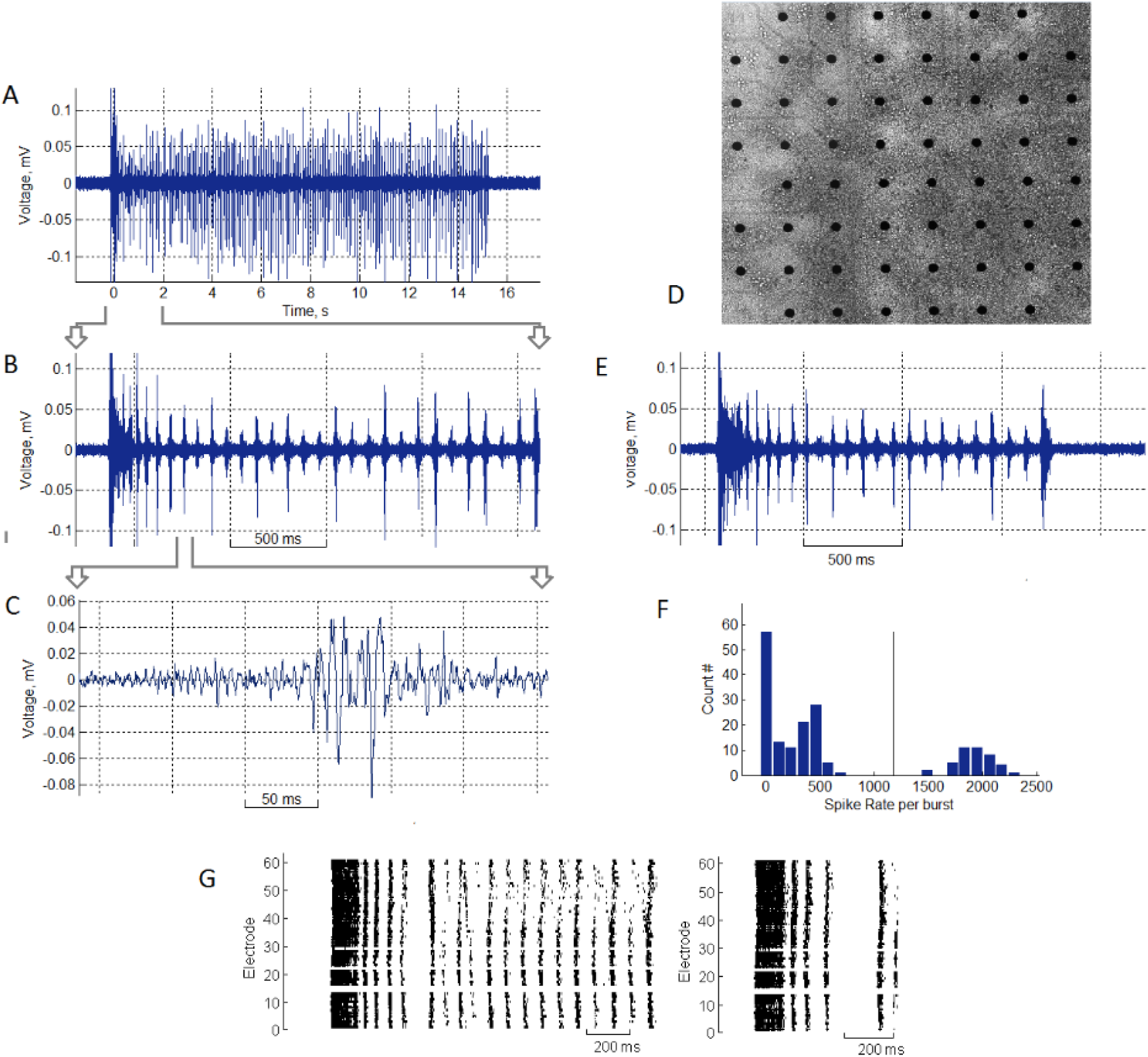
Long superburst activity in hippocampal culture recorded by the microelectrode array at DIV 35. (A) Electrophysiological signal of spikes within a long superburst recorded from a single electrode. (B) Long superburst at a 2 s timescale and (C) a 200 ms timescale. (D) Dissociated hippocampal neurons grown on a microelectrode array (DIV 35). (E) Regular superburst activity with a duration of 1-2 seconds. (F) Initiation burst and subsequent small bursts can be separated by clustering the numbers of spikes per burst. The threshold (the vertical line) was identified using K-means clustering. (G) Raster of a long superburst (left) and a regular superburst (right).

Phase-contrast images of cultures were taken weekly to record the status of the culture using a Leica DMIL HC (Germany) inverted microscope with 10x/0.2 Ph1 objectives. Experiments were conducted when the cultures had been grown for 3-5 weeks *in vitro*.

### 2.2 Electrophysiological methods

Extracellular potentials were collected using 59 planar TiN electrodes integrated into the USB-MEA-120 system (Multichannel system, Germany). The microelectrode arrays (MEA) had 59 electrodes (8x8 grid) with diameter of 30 μm and spaced 200 μm apart (Fig 1 A). Data were recorded simultaneously from 59 channels at a sampling rate of 20 kHz/channel. All signal analysis and statistics were performed using the custom-made software Meaman in Matlab (Mathworks, USA).

### 2.3 Spike detection

The detection of recorded spikes (Fig 1 B) was implemented using threshold calculation:

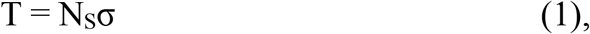

where *σ* = median (|*x*| / 0.6745), which was the estimate of the median normalized to standard deviation of a signal with no spikes (see [50] for more details), x is the band-pass-filtered (0.3-8 KHz) signal, and NS is the spike detection coefficient, which was set to 8. The amplitudes of detected spikes were in the range of 20–200 μV. The minimal interspike interval was set to be 1 ms to avoid the overlapping of neighbouring spikes.

### 2.4 Burst detection

The burst detection method was described in detail in our previous paper [31]. Briefly, we estimated the total spiking rate characteristic, TSR(t), as the number of spikes from all electrodes within each 5 ms time bin. Fast appearance of a large number of spikes over all electrodes in a small (2 ms) time bin was used as the criterion for burst appearance. Threshold detection was applied to estimate burst beginning and ending points. The burst threshold was set to T_Burst_ = 0.2 × σ_TSR_, where σ_TSR_ is the standard deviation of TSR(t).

The initiation time of the burst was defined as the burst start time, where TSR(t) was above the threshold. Next, the initiation time was adjusted to the first spike from all electrodes after a supra-threshold time. Finally, the time point at which TSR crossed the threshold after the burst start time was defined as the burst end time.

### 2.5 Superburst detection

Superburst and long superburst in the electrical activity were detected using the method described previously [51]. First, we defined a Gaussian function with an effective width equal to 50 s. Next, that function was iteratively moved from the beginning of the recording to the end using a 10 ms time step, while cross-correlation of the function with the TSR was calculated for each step. The resulting cross-correlation indicated how much of the synchronized activity (bursts) was recorded in each 10 s window. To detect superburst in the spiking activity, we applied a threshold detection algorithm in which the threshold was estimated as the superburst detection accuracy coefficient multiplied by the standard deviation of the calculated cross-correlation. The superburst detection accuracy coefficient was estimated empirically and was equal to 0.4. All time points that crossed the threshold were defined as the beginnings and the endings of the superburst, respectively [51].

### 2.6 Burst classification

Superburst consisted of initiation bursts lasting 50-100 ms and short small bursts lasting 30-50 ms. The total number of spikes within initiation bursts was in the range of 1000-3000 spikes, while small bursts each contained 10-500 spikes (Fig 1 F). These two types of bursts were identified using a K-means clustering algorithm.

To represent spatio-temporal properties of all patterns within small bursts, we analysed activation patterns consisting of first spike timings of the bursts. The first spike timing was averaged for each electrode and each small burst. Then, the values from all 60 electrodes were colour coded and plotted on an image using cubic convolution interpolation. That image represented a gradient map of burst activation profile.

To represent spatial properties of spike propagation during burst activation, we introduced a vector field map of activation timings in the culture. For each MEA electrodes we calculated a vector whose direction represented the activation-timing gradient around an area of 3 electrodes. The resulting vector field resembled a colour-coded activation pattern. We defined this spatial representation of the activation pattern as a *dynamical pattern*.

To identify motifs of activation patterns in small bursts of all long superbursts, we applied the K-means clustering method. The activation patterns for each small burst consisted of the first spike timings for each of the 59 electrodes of the MEA. This method required a number of clusters to be estimated. First, we estimated two clusters and evaluated cluster separation by calculating the Davies–Bouldin (DB) index [32]. The DB index estimated the ratio between internal cluster distance and distance between clusters. Then, the clustering procedure was repeated for various numbers of clusters (2, 3…30), and the DB index was estimated. We found that the minimum DB index value among all tested clusters numbers was 2 or 3, indicating that the activation patterns were optimally clustered into 2 or 3 motifs of small bursts. Small values of DB-index corresponded to compact clusters whose centers were far away from each other. DB values in the range of 0 to 1 indicated “robust” clustering. We also tested the motif separation by expectation–maximization algorithm (EM clustering). To represent clustering evaluation, we plotted two Principal Component coefficients for each activation pattern and highlighted the estimated clusters in different colours. As in the previous case, we used the DB index to evaluate the optimal number of clusters with EM clustering. This method applied to 3 Principal Component coefficients divided data into the clusters more accurately and, hence, was used in further analysis.

## 3. Results

First, we analysed spontaneous activity of the hippocampal cultures. We obtained complex bursting patterns similar to those reported previously in cortical cultures [26]. An example of the spikes recorded from single electrode within a small burst is shown in Fig 1 C. After 3-4 weeks *in vitro*, we obtained the activity described as superburst (Fig 1 E). A typical superburst consisted of a sequence of 3-20 small bursts of 50-100 ms in duration and a 50-150 ms interburst interval. During the period of 30-40 DIV, the cultures generated long superbursts that were similar to regular superburst but lasted longer, 10-30 seconds, and consisted of hundreds of regular small bursts. In summary, we analyzed 11 cultures from 3 plating experiments and observed long superbursts in 8 cultures. We found that 6 cultures generated more than 6 long superbursts at least during 20 minutes which were included in the statistical analysis. In other cultures we observed no more than 2 long superbursts. The signals from a single electrode during the long superburst on timescales of 15 and 2 seconds are illustrated in Figs 1 A and B. Raster plots of the spiking activity recorded from all 59 electrodes during the superburst and long superburst are shown in Fig 1G. Each point on the raster represents the spike occurrence time for a particular electrode. The long superburst were composed of relatively long initiation bursts (50-100 ms) followed by shorter bursts, i.e., the small bursts. The initiation bursts and the small bursts can be easily identified by K-means clustering (Fig 1 F) using burst firing rate features (see Methods).

Next, we estimated the frequencies of the small bursts, the initiating bursts and intervals between the long superburst were excluded from analysis. *Interburst peak intervals* (IBPIs) were calculated as the time interval between adjacent peaks from the *total spike rate* (TSR) diagram (see Methods) of detected small bursts (Fig 2 A). Each IBI corresponded to the *instantaneous frequency* (IF) of each pair of the small bursts. Note that the IFs were not normally distributed (Kolmogorov-Smirnov test, p<0.01) and for each culture the median value of the IF was estimated. Typical example of frequency distribution for a small burst sequence in one culture is shown in Fig 2 B. Note also that the IBPIs were highly stable, having median frequency at 9.8 Hz. Less than 5% of the bursts were in the range of 15-30 Hz. Then the medians were averaged for the all cultures and the IF was equal to 11.2 ± 1.5 Hz (mean ± standard deviation, n=6 cultures). Average histogram of the IFs for the cultures with long superbursts is illustrated in Fig. 2 D. Most of the IFs were concentrated in range from 8 to 15 Hz.

**Fig 2.**
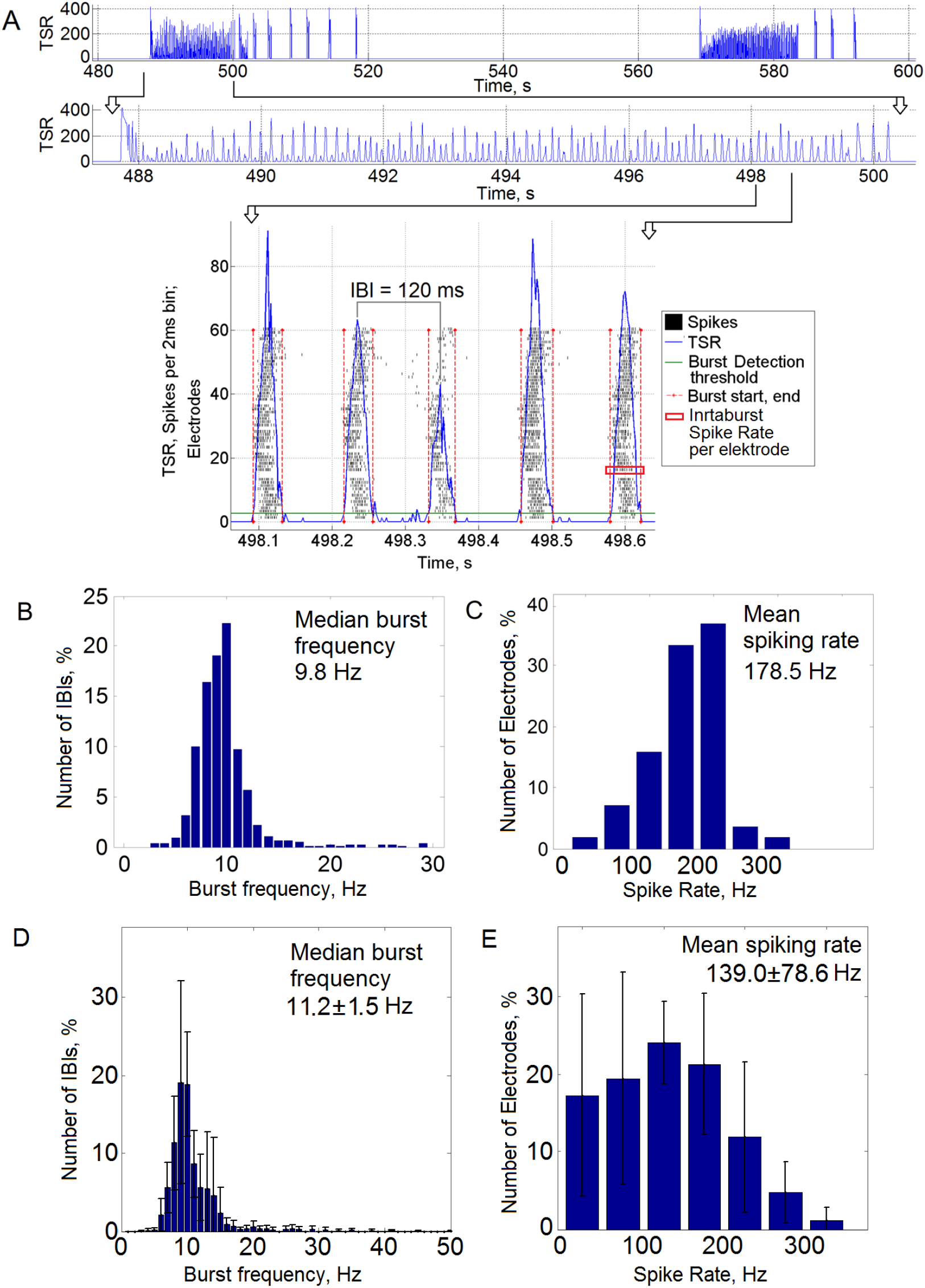
The sequence of small bursts in a long superburst had a stable rhythmic structure with a frequency similar to the hippocampal rhythmic activity. (A) Example of long superburst activity recorded on MEA and a fragment of 5 detected small bursts (bottom). The green horizontal line represents the burst detection threshold, and the red vertical lines represent the burst initiation and end time points. (B) Distribution of burst frequencies in one culture. The median burst frequency was 9.8 Hz. (C) Distribution of spiking rate frequency from the electrodes in the small bursts in one culture. The mean spiking rate was 178.5 Hz. (D) Distribution of burst frequencies (n=6 cultures). The median burst frequency was 11.2 ± 1.5 Hz (mean ± s.d., n=6 cultures). (E) Distribution of spiking rate frequency from the electrodes in the small bursts (n = 6 cultures). The spiking rate was 139.0 ± 78.6 Hz (mean±s.d.).

We also estimated the mean spiking frequency for each electrode during small bursts only in intra-burst periods (Fig. 2 A, red rectangle marked period in sample burst). For the raster presented in Figure 1 A the most of the electrodes had spiking frequency in range 100-300 Hz, and mean frequency was equal to 178.5 Hz (Fig 2 C). On average the spiking frequency per electrode was 139.0 ± 78.6 Hz (mean ± standard deviation, n=6 cultures, 264 active electrodes) (Fig. 2, E).

Next, we analysed spiking patterns in sequences of small bursts within long superburst recorded for 30 minutes. The set of the first spikes in the burst recorded from all electrodes was considered as activation pattern [31]. To investigate different motifs (clusters) of activation patterns, we applied the EM clustering algorithm for 3 Principal Component features. This method required the estimation of a number of clusters. First, using two clusters, we evaluated the cluster separation by calculating the Davies–Bouldin (DB) index (see Methods). Then, the clustering procedure was repeated for various numbers of clusters (2, 3…30), and the DB index was estimated (Fig 3 A). We found that the minimum DB index value among all tested numbers of clusters was 3, indicating that the activation patterns were optimally clustered into 3 motifs in the presented raster. The activation patterns and the profiles of the spiking patterns in the bursts from separate clusters (motifs) represented different sequences of spike occurrence, i. e., different dynamics of the spike propagation (Fig 3 B). Note that the difference in the first spike time sequence in the pattern can be observed visually from Fig 3. Examples of the raster plots for all of the small bursts from all 3 motifs are shown in Figs 3 C, D and E. To investigate the spatio-temporal properties of all patterns within each motif, we averaged the activation patterns and calculated the dynamical patterns (see Methods (Figs 3 F, G, H)). The first spike timings from all 59 electrodes were colour coded and plotted on the image created using cubic convolution interpolation. Surprisingly, we found that the burst activation was implemented in the form of wave-like spike propagation dynamics with a wide wave front. Arrows represent the gradient of activation time, i.e., the mean direction of spike propagation during burst initiation across each electrode. Eventually, the patterns were organized with a uniform direction in space. Remarkably, motifs #1 and #2 presented similar directions of the activation pattern, from the upper electrodes to the bottom of the MEA, whereas motif #3 presented the opposite direction.

**Fig 3.**
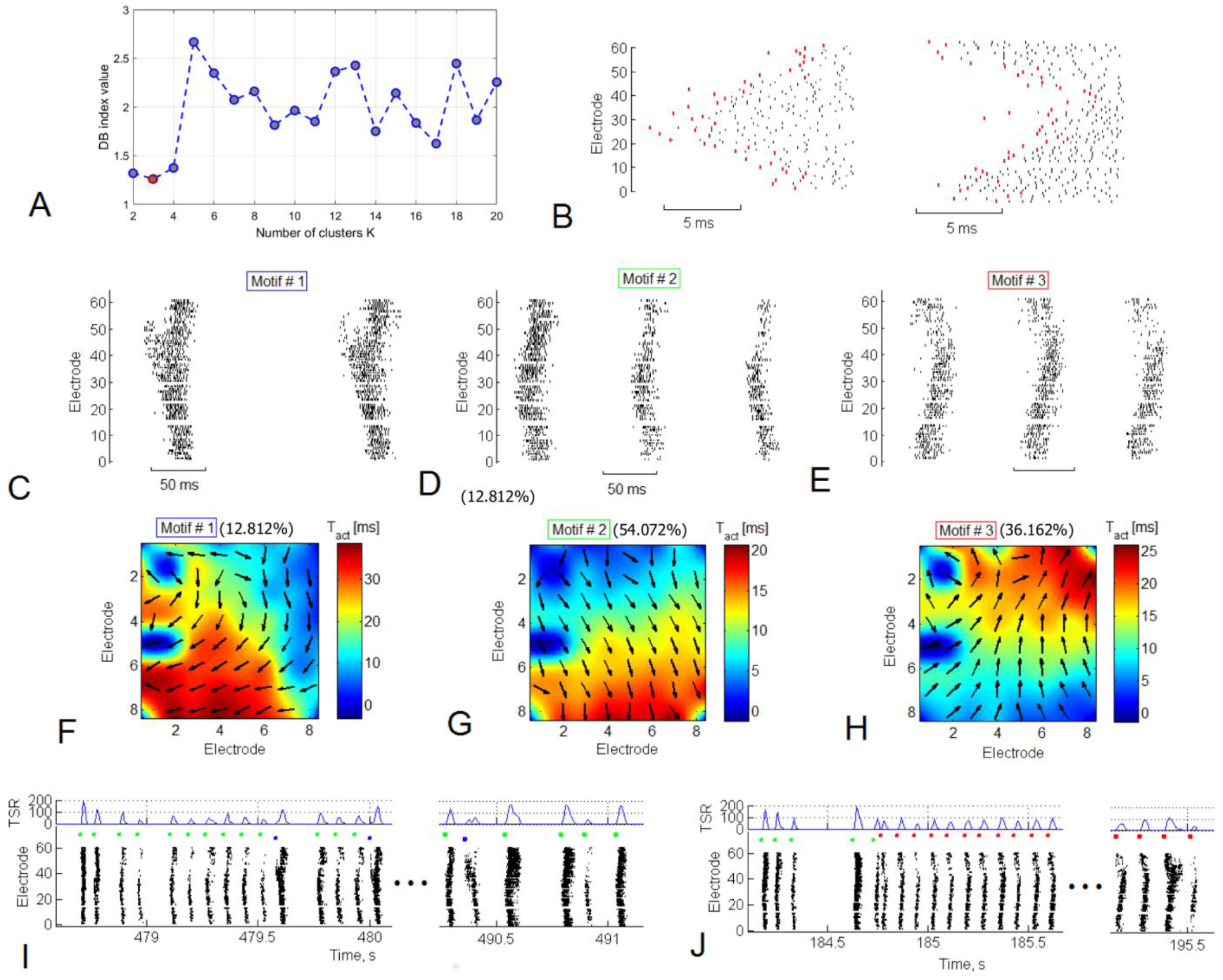
(A) Small bursts from several long superburst can be clustered into spatio-temporal patterns. The DB index (see Methods) showed a minimum of 3 clusters, indicating that there are 3 most dissimilar groups (motifs) of small bursts according to the activation spiking pattern. (B) Examples of the bursts from two motifs. Red dots indicate activation pattern - first spike timings for each electrode. Examples of the bursts from 3 motifs (C, D, E) and spatial representation of *dynamical patterns* (F, G, H), respectively. Motif #1 was observed in 10.6 *%* of small bursts, motif #2 in 56.4% and motif #3 in 32.9 *%* of all small bursts in superburst. Examples of two types of long superburst (I, J) that were composed of the 3 motifs. The left long superburst consisted of motif #1 bursts (blue markers) and motif #2 bursts (green markers) (F). The second type of long superburst consisted of bursts of motif #3 (red markers, J).

Next, we analysed the sequence in which the motifs appeared in the structure of the long superburst. Some of the long superburst (Fig 3 I) were composed of bursts of motifs #1 and #2, whereas other long superburst in the same recording (Fig 3 J) were composed mostly of bursts of motif #3.

Note that the firing rate in the burst sequence (Fig 3 J, TSR - total spiking rate) was quite variable but that the sequence of the first spike timings, i.e., the activation patterns, of the bursts remained largely unchanged.

Furthermore, we verified the clustering results for activation patterns using EM clustering for two *principal components* (PC), three PC and K-means clustering (see Methods). We applied all three methods to one data set: varied the number of clusters and estimated clustering using DB index as in Fig. 3 A. We found that for K-means and EM clustering with 2 PC the DB index had minimum at 2 clusters (Fig. 4 C and F), while DB index for EM with 3 PC had minimum at 6 clusters (Fig. 4, J). Interestingly, whole activation patterns (spike timing over 59 electrodes) reduced to 2 *principal components* were clustered into just two motifs, whereas the patterns reduced to 3 PC - into 6 motifs. To visualize clustering results we illustrated all patterns by a color scale on 2 PC plot for K-means and EM 2PC (Fig. 4 A and D) and on 3 PC plot for EM 3PC (Fig. 4 H). Note that even without clustering the patterns in 3 PC space can be visually identified as 6 clustered set, while K-means and EM 2PC could identify only 2 clusters. However, among 6 motifs one can note 2 motifs found by other algorithms (Fig. 4 B, E, I). On average, the optimal number of motifs (minimum DB index) found by the EM clustering for 2 PC (Fig. 4, G) and EM for 3 PC (Fig. 4, K) displayed clear minimum in 2 clusters (n=6). Interestingly, in all cases visual inspection of the clustering demonstrated two major motifs associated with global spike propagation pathways across MEA. To emphasize this we applied the following analysis.

**Fig 4.**
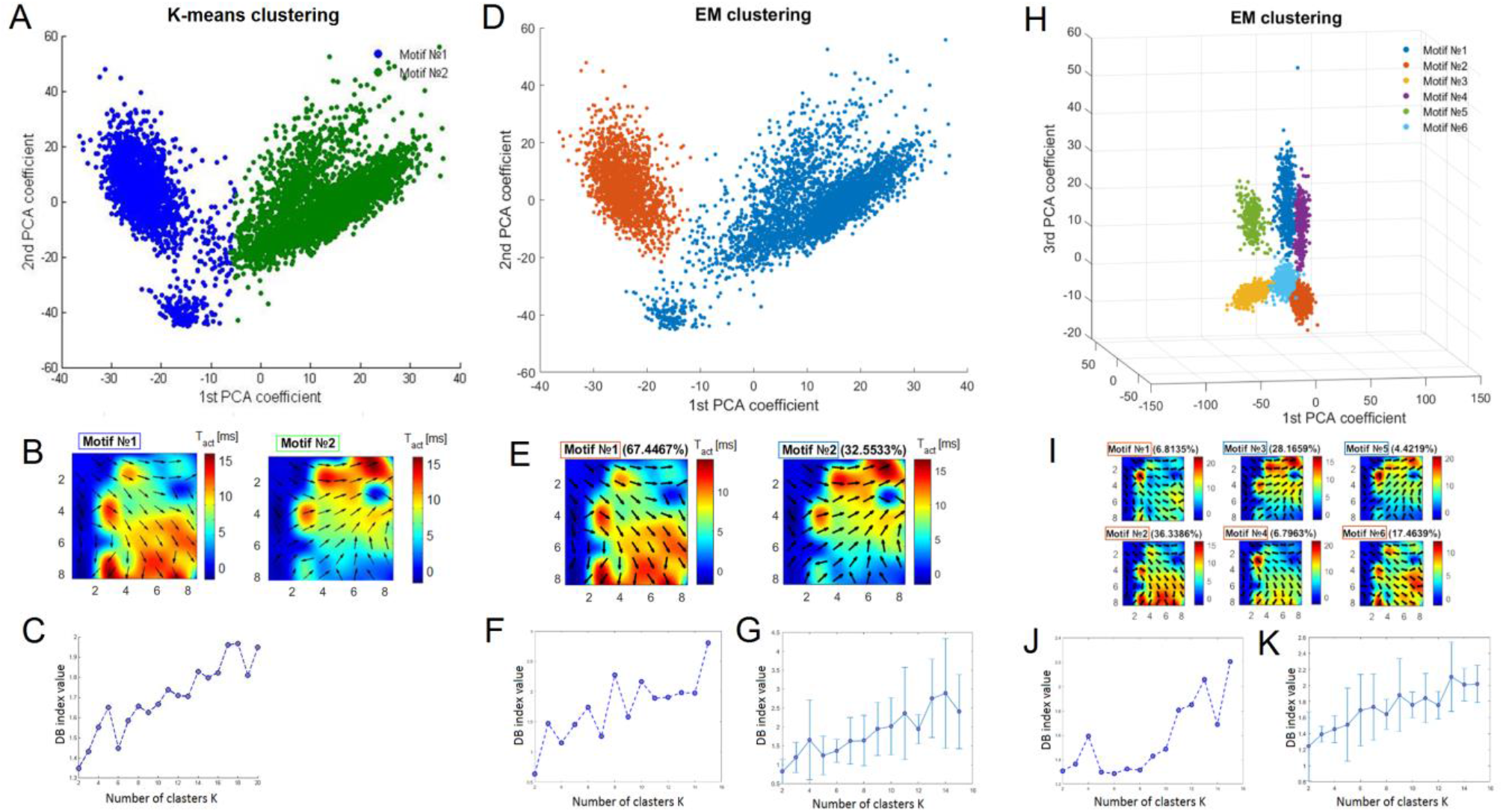
Clustering of the spiking patterns of small bursts in long superburst. (A) Clustering of the activation patterns with K-means revealed two motifs (green and blue dots) plotted on principal component analysis (PCA) coefficients. (B) Dynamical patterns as average activation patterns of two motifs found with K-means clustering. Colour represents the average first spike timing of the bursts. (C) Dependence of the Davies–Bouldin (DB) index on the number of clusters, as estimated by K-means. (D) Clustering of the same spiking patterns using EM clustering applied to two principal components. (E) Dynamical patterns of motifs found with EM clustering. (F) Dependence of the DB index on the number of clusters, as estimated by EM clustering and (G) its average (mean±s.d., n=6 cultures). (H) EM clustering of the spiking patterns applied to three principal components estimated 6 motifs. (I) Dynamical patterns of 6 motifs estimated by EM with 3 principal components. (J) DB index dependence on the number of clusters, estimated by EM clustering with 3 principal components and (K) its average (mean±s.d., n=6 cultures).

The activation patterns can be characterized by one major direction of spike propagation pathway. For each pattern we estimated the angle of *major direction* by averaging all 59 vectors (Fig. 5 C). Next, the clustering of the major direction angles revealed two major *direction motifs* in the raster (Fig. 5 D). The DB index value was equal to 0.08 which represent robustly separable two clusters, which also can be seen on a major direction angles histogram (Fig. 5, F). Mean of the angles from two motifs were equal to 29° and 302° and the difference between them was statistically significant (t-test, p<0.01). Interestingly, clustering of the activation patterns composed of spike timings (EM 2 PC, Fig. 4, E) and major directions angle showed similar dynamical patterns (Fig. 5, G). Average DB index of the major direction angle clustering showed that in all cultures (n=6 cultures) with long superbursts the patterns clearly clustered into two major directions (Fig. 5, E). Minimum value of the DB index in the cluster estimation was equal to 0.33±0.27 (mean and s.d.). Note, that DB index value below 1 indicated well separated clusters when the intercluster distance was higher than the intra-cluster volume.

**Fig 5.**
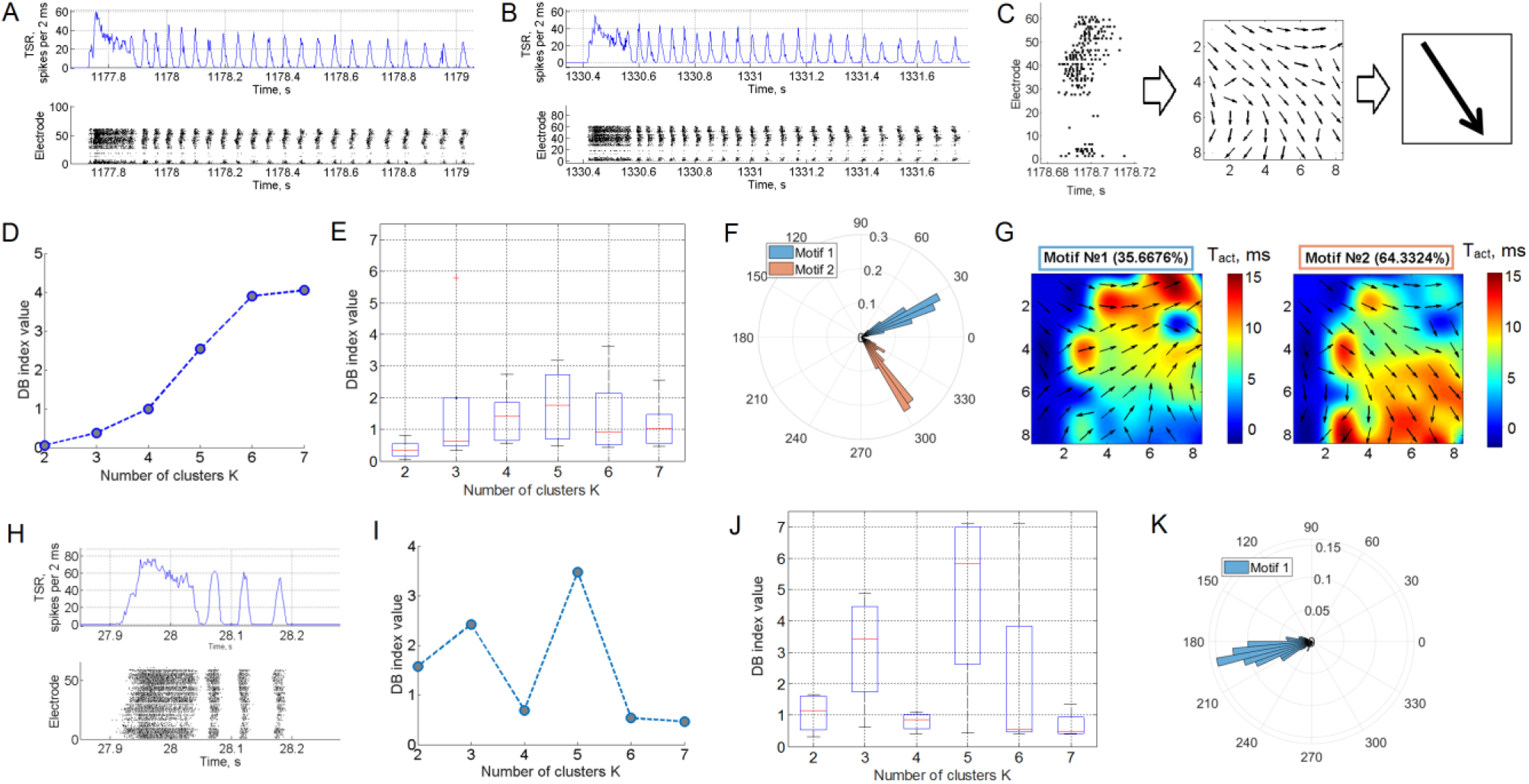
Bidirectional spike propagation pathways form two types of long superburst. Rasters of two types (A and B) of the long Superburst obtained in one recording (bottom) and its TSR diagram (top, see methods). (C) Schematic view of major direction in spike propagation estimation. (D) DB index dependence on cluster number in clusterization of major directions in the long superburst for example raster and (E, boxplot) for 6 cultures. (F) Histogram of major directions of the bursts from the culture with long superbursts. Two colored clusters of the histogram represented two motifs. (G) Spatial representation of dynamical patterns of motif #1and motif #2 indicated different spike propagation pathways after clusterization of major directions. (H) Raster of superburst (bottom) and TSR diagram of the superburst (top, see Methods). The superburst composed of an initial burst (100-150 ms) followed by a subsequence of 3-10 small bursts. (I) DB index dependence on cluster number in clusterization of all major directions of the small bursts in superburst. (J) Boxplot of DB index of the small bursts in superbursts (L) (n=5). (K) Histogram of major directions of the bursts from culture with superbursts.

To test whether such well-defined bidirectional dynamics is a unique feature of long superburst, we shifted from analysing long superburst to analysing regular superburst for the presence of stable spike-propagation pathways. Such superburst are also composed of an initial burst (100-150 ms) followed by subsequent 3-10 small bursts (Fig 5 H). The profile of the small bursts within such superburst was less clearly organized than we observed for the long superburst. In many cases, the small bursts cannot be clearly separated due to the high variability of activity in development. We found that for this example 2 clusters could not be estimated correctly because one cluster contained more than 95% of all patterns. Also, the DB index characteristic was not monotonic (Fig. 5, I) in contrast to long superburst clustering (Fig. 5, D) suggested the absence of the motifs. Indeed, histogram of all major directions from this raster plot (Fig 5 K) indicated the existence of one cluster (e.g. motif) with average angle of 192°. DB index characteristics averaged over 5 rasters (n=5 cultures) did not indicate any clustered structure. Note, that for 2 clusters average DB index was equal to 1.06±0.65 (mean and s.d., excluding one sample with <5% in one motif) which was significantly different from the mean DB in the long superbursts 0.33±0.27 (n=6 cultures, t-test, p<0.05). In average, the cluster analysis of the short bursts did not reveal a clear minimum for the DB index in 2 clusters (Fig 5 J, boxplot, red lines - median values) in contrast to the recordings with long superburst (n=6 cultures) (Fig 5 E). Thus, the patterns in the long superbursts were clustered into two motifs associated with significantly different spike propagation directions, while regular superbursts did not show such a feature. Note that mean DB index for the long superbursts 0.33±0.27 was lower than 1. Such low values indicated two clusters without inter-cluster overlapping [52] and, hence, can be treated as statistical evidence of error-free clustering.

Then we analyzed the reproducibility of motif appearance in long superburst sequence. Motifs in the long superburst were represented using raster plots (Fig 6 A). The vertical black lines in the plot represent the motifs in the sequence of the small bursts in all long superburst. We applied EM clustering to the long superburst based on their motif frequency. This analysis identified two clusters associated with the two types of the long superbursts, which are marked in blue and pink color in the motif raster (Fig 6 A, top). Note, that each type appeared randomly in the sequence. Surprisingly, a switch probability between the two types of the long superbursts was equal to 50% ± 13% (n=6 cultures), indicating random nature of such activity in macroscopic timescale. Each long superburst was composed of 100-150 small bursts, which can be clearly associated with a certain major angle of the motif (Fig 6 B). Surprisingly, the first type in the presented example was composed mostly of small bursts of motif #1 (96.9% of all small bursts) and partially of motif #2 (3.1%). This fact indicated the presence of stable and directed spatio-temporal patterns of spike propagation pathways during the long superburst activity. In contrast, the other type of long superburst was mostly associated with motif #2 (motif #1 −18%, motif #2 −82%), representing different direction of spike propagation. Figure 6 C shows dynamical patterns of two estimated motifs where these major directions can be clearly seen. In average, each motif appearance probability within its own type of the long superburst was equal to 91.5% ± 4.7% (n=6 cultures). Such remarkably high appearance of the motif in the long superbursts clearly demonstrated the presence of stable functional structure of the network and, hence, reproducible dynamics.

**Fig 6.**
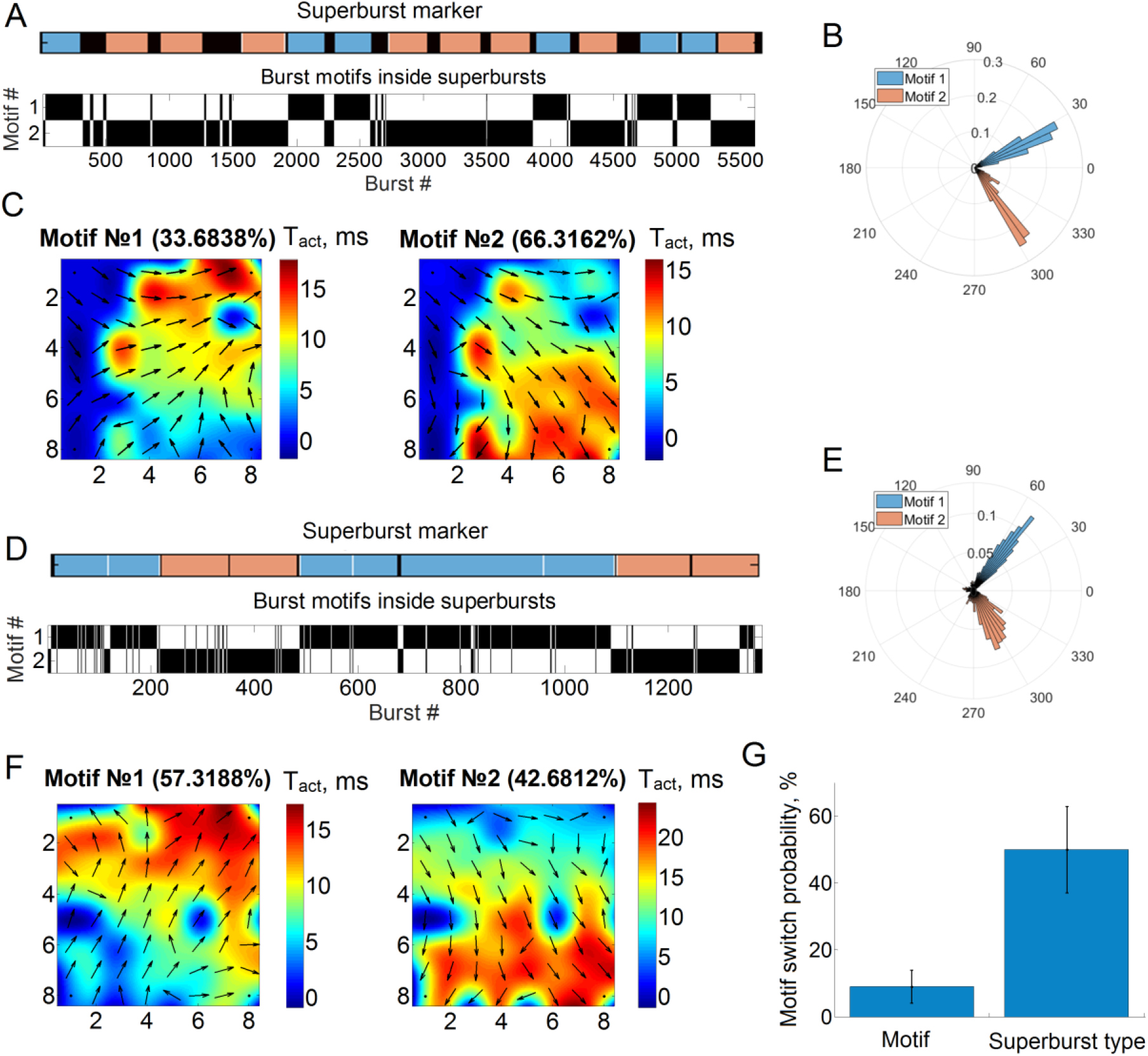
Spike propagation pathway switch during long superburst. (A) Motif appearance within each superburst. Blue and pink bars mark two types of long superbursts in one recording. Black vertical lines illustrate bursts of a particular motif type inside each superburst. Note that one superburst type (blue) was associated with motifs #1, and the other one (pink) was associated with motif #2. (B) Histogram of major directions for small bursts from culture with long superbursts. Two colored clusters of the histogram represented two motifs after clusterization, which appeared in (A). (C) Motif dynamical patterns #1 and #2 with respect to B. Note that in all cases, the motifs from different types of superburst have clearly different activation gradients (arrow directions) and major directions (B). The other culture showed similar principal results using the same analysis: two motifs were associated with two types of long superbursts (D), bursts from the motifs had significantly different major directions (E, DB index <0.1) and spatio-temporal activation patterns (F). (G) Switch probability of burst patterns during a long superburst (motif) and between subsequent long superbursts (superburst type).

To quantify the stability of the motifs in the small bursts sequence, we also measured the probability of switching between the motif types in the whole raster without considering long supeburst indexing (Fig 6 A). We found that the switch probability was quite low −9% ± 5% (n=6), which shows that motif switched quite rarely Thus, spontaneous bursting activity employed two basic spike propagation pathways (types of motifs) that were activated and sustained during long superburst (10-20 sec). Examples of similar bidirectional activity of the bursts from another culture is shown in Figs 5 D, E, F. Notably, motifs from different long superburst types eventually show significantly different (DB index directions of spike pattern propagation using major direction measure or dynamical pattern representation. For each motif we estimated mean major angle and the difference between two mean angles. In average, the difference between them were equal to 95° ± 31.1° (mean± s.d. n=6 cultures). Surprisingly, such almost perpendicular spike propagation pathways were self-replicated in 6 cultures spontaneously.

## 4. Discussion

In this study, we discovered that neuronal networks formed in hippocampal cultures in their mature state (30 DIV and older) generated specific network activity with surprisingly long sequences of bursts, i.e., long superburst of up to hundreds of constituent bursts, with highly regular spiking patterns. In previous studies, superburst activity was reported with much shorter duration (up to ten bursts) [26]. In our experiments, we also observed similar activity, but in addition, more than 70% of the cultures (8 out of 11) at DIV 30-35 began to spontaneously generate long superburst with durations of up to hundreds of seconds. We found 6 cultures out of 11 generating more than 6 long superburst at least during 30 minutes. The other cultures generated less than 2 long superbursts with a regular superburst in the background. A precise biophysical mechanisms of the long superburst are still largely unknown we suggest that for matured cultures (DIV from 35) spontaneously organized in the well balanced excitation-inhibition networks where cycling dynamics represents homeostatic (“natural mode”) pattern “sustaining” the functional connectivity.

This long superbursting activity persisted for several days. Each superburst consisted of an initial burst with the highest firing rate, followed by a subsequence of small bursts with 50-100 ms duration and a relatively stable 100-200 ms interburst interval (Fig 1). This interval corresponded to the bursting frequency 11.2 ± 1.5 Hz (n=5) (Fig 2) which represented a hippocampal rhythmic activity [49]. Such periodic bioelectrical activity in a form of theta oscillations in cultures can be considered a fundamental feature of hippocampal network formation, which has been widely investigated *in vivo* and in slices *in vitro* [7–11]. In hippocampal neural networks the theta frequency range was defined at 4-10 Hz and the beta range at 10-30 Hz [53, 54]. Other authors define the theta range at 4-12 Hz and beta at 12-30 Hz [55]. Our analysis suggests the features of both rhythms and indicate a fundamental feature of hippocampal network to generate theta rhythm for behavioral (active motor behavior [49]), memory tasks and also by complex electrophysiological signatures of sleep state [14] in theta and beta range. Low-frequency synchronized firing in cortical cultures was associated with classical sleep signatures [14]. The lack of external stimuli during the neural network development could trigger such stable low-frequency spiking activity in cortical and hippocampal cultures and indicate functional state of sleep [14].

The spiking rate in the small bursts was found to be 139.0 ± 78.6 Hz (n=6), which we suggest may be associated with high-frequency oscillations and sharp wave-associated ripples in the hippocampus (100–250 Hz) [7, 10] or fast gamma oscillations (90–140 Hz) [56]. Such a high spiking frequency was also observed in regular superburst and hence cannot be directly associated with the unique characteristics of the observed long superburst. However, the interplay between the dynamics of small bursts and the generation of high-frequency spiking plays important functional roles *in vivo*. High-frequency oscillations have been found to be modulated by slow theta activity in isolated rat hippocampus [57] and *in vivo* [58]. Interestingly, coupling of the theta and high-frequency oscillation has been observed during REM, slow-wave sleep and immobility behaviour, whereas ripples have been associated with memory consolidation [10].

We suggest that such unique and rare activity appeared in cultures in which a specific balance of morphology and cell density developed spontaneously from the specific initial conditions due to mechanisms of selforganisation. It has been shown that the cell density of neurons surviving to DIV 30 decreased dramatically from 2500 cells/mm^2^ to 150 cells/mm^2^ [36]. In another study, the cell density was decreased from 5000 cells/mm^2^ 2500 cells/mm^2^ after two weeks of culture development [46]. Hippocampal cultures (E18) with high density of 10^6^ cell/mm^2^ in DMEM medium generated superbursts with long duration in range of 46-91 seconds with relatively low interbursts frequency of 0.4 - 4.6 Hz [43], which was associated with epileptiform activity[61]. Superbursts in the rat hippocampal cultures E17–E18 with high density of 7800 cells/mm^2^ were also observed in [42, 60], but detailed activity analysis was not presented. Superbursts with 10 Hz frequency were observed only in conditions of increased cAMP in 600 cell/mm^2^ cellular density [63] or with additional inhibitory cells dissociated from striatum [47] in the cultures with 1000 cell/mm^2^. In our cultures initial cellular density was 15000–20000 per mm^2^. The plated culture contained about 250000 cells and formed 4-5 layers. To our knowledge it was the first time that the spiking activity was observed in such plating conditions at that stage of culture development. Thus we suggest that the plating density of the cultures was the major factor responsible for the appearance of such activity.

Note that the culture medium was changed every 2 days in our experiments, while in other studies the medium was changed once or twice a week [15, 26]. In this aspect we suggest that frequent change of small amounts of the medium minimized physiological stress and sustained homeostatic balance during culture development which could affect on cultured network stable activity. It has also been shown that the densities of glutamatergic and GABAergic synaptic terminals increased during first 3 weeks *in vitro* and then saturated by DIV 30-35 [63]. In cortical cultures [63] and in hippocampal cultures [64, 65], the ratio of glutamatergic to GABAergic receptors in synapses and somata during development showed a similar tendency to that observed *in vivo*. We suggest that the ratio of inhibitory and excitatory cells and the cell density formed by DIV 30 in our experiments were also important components for the development of the *in vitro* neural networks with such remarkable features of activity and will be investigated in detail in further studies.

Many studies have shown that theta rhythmic activity in the hippocampus can be induced and modulated by external signals originating from the entorhinal cortex. Our results suggest that the spiking activity in the range of the hippocampal theta rhythm can be generated in isolated networks of hippocampal cells without any stimulation applied under stable homeostatic conditions. Further studies using immunohistochemistry will reveal key features of such culture conditions.

Important to note, that in cortical cultures the superbursting activity did not demonstrate such long and stable rhythmic activity [14, 22, 26]. Reverberation of activity in a form of periodic synchronized bursts on a time scale of hundreds of milliseconds emerged or modulated only under suppression of inhibitory synaptic transmission [18, 48]. Superbursting activity in dense cortical cultures (1000-5000 cells/mm^2^) of rats and mice consisted of sequence of the bursts in range from 0,25 to 1,25 Hz [26,46,67–69]. Such activity was associated with epileptic seizures *in vitro* [61]. In rat hippocampal slices similar epileptiform bursts had high-amplitude (1 mV) and low-repetition frequency (0.5 – 1.5 Hz), whereas theta oscillations had a low-amplitude (0.5 mV) and high frequency (5–14 Hz) [70, 71]. It was also shown that in cortical cultures of postnatal neurons with high density (4000 cells/mm^2^) in some cases the superbursts were generated with frequency of 10 Hz, and could be induced by GABA synaptic transmission blockers [18]. Therefore, the observed result of the bursting activity in theta and delta range could be unique to the hippocampal cultures.

Note that the recording area of the MEA (1.6 x 1.6 mm) was in the centre of the circular culture and had an approximate diameter of 4-5 mm. These spatio-temporal patterns may be part of a global cycling activity with highly stable specific features of the hippocampus *in vivo -* theta rhythmic oscillations. The cycling pattern of the bursting activity might be triggered by pacemaker neurons [72] or may be self-organized in spiral wave dynamics. Similar spike propagation patterns were observed in cortical cultures with chemically mediated inhibition in the network [13]. Disinhibition of GABAa synaptic transmission by *biccucline* induced episodes of seizures composed of stable short burst subsequence with 2-3 sec of interburst intervals. Further studies using high-density MEA systems, or fast CCD cameras for calcium imaging [61], to observe the activity of the whole culture will address such questions.

Analysing the profile of the spiking patterns in the long superburst, we found that they also become well organized, containing a small number (2-4) of basic motifs, in contrast to regular superburst activity (Figures 4 and 5). Using clustering methods we found that the patterns of first spike timings in the bursts have two clusters (motifs). We used DB index measure to estimate clustering efficacy. We found that DB index for long superbursts was 0.33±0.27 (mean, s.d.). DB value in that range implies that significant parts of two clusters do not overlap (see Methods), thereby leading to robust clustering.

Such motifs defined the presence of two basic types of spike propagation directions in the burst activation pattern. These two “functional” directions further defined the activity in the form of wave-like bidirectional firing patterns repeated from burst to burst. The stability of the motif appearance within single long superbursts were equal to 91.5% ± 4.7% (n=6 cultures). Such remarkable stable pattern reentry clearly demonstrates stable functional structure of the network. Considering spike timings variability in the culture and clusterization inaccuracy such motif uniqueness would be even closer to 100%. Notably, the angle between two major spike propagation pathways of the small bursts activation patterns were equal to 95° ± 31.1° (mean± s.d. n=6 cultures) (Fig. 6 B, E). Such almost perpendicular activity propagation pathways were selforganized in 6 cultures spontaneously and were remarkably stable during such rhythmic bursting. The long superbursts were also clearly clustered into two types according to the motif appearance in small bursts subsequence (Fig. 6, A). Each motif appeared mostly within its own type of the long superburst with high probability 91.5% ± 4.7% (n=6 cultures) indicated that during each long superburst only one of two motifs was generated. We suppose that on a timescale of milliseconds (activation patterns), seconds (small bursts) and tens of seconds (long superbursts) the dynamics of the neural network was quite stable and reproducible. Interestingly, on a timescale of minutes where several long superburst can be observed in the activity pattern switched from one motif to another with 50% ± 13% (n=6 cultures) chance. The switch appeared mostly from the first small burst in the sequence. We suggest that the first initiation burst with higher duration and firing rate in the long superburst sequence determined that motif. Therefore, the spiking patterns with unique orientation of activity propagation demonstrated an interplay of two complex dynamical states in the network with stochastic switch on a several minutes timescale which initiated rhythmical activity with precise activity pattern on a timescale of seconds and milliseconds. Such conclusion complements the results of regular superburst study [22] and can be further extended to the modeling of brain dynamics development. The results suggest that hippocampal neuronal cultures can demonstrate the activity with the features similar to *in vivo* conditions. Further study of plating protocols and neuroengineering methods mimicking realistic hippocampal tissue conditions can uncover key factors of functional structure development in the neural networks.

Notably, such well-organized global dynamics of spontaneous culture networks can be encoded by a binary sequence and represented, in fact, as a telegraphic signal conveying information about the functional state of the system. We sincerely believe that such stability and reproducibility of the network states will further permit the control of the switching between the states in mature cultures and that such culture will be useful in the design of living networks with definite functional properties in hybrid information processing systems (neurally controlled robots, “brain-on-chip”, etc.).

## Acknowledgements

This research was supported by the Russian Science Foundation (No 16-19-00144).

## References

1. Vida I, Bartos M, Jonas P. Shunting inhibition improves robustness of gamma oscillations in hippocampal interneuron networks by homogenizing firing rates. Neuron. 2006;49: 107–117. doi:10.1016/j.neuron.2005.11.036

2. Brunel N. What Determines the Frequency of Fast Network Oscillations With Irregular Neural Discharges? I. Synaptic Dynamics and Excitation-Inhibition Balance. J Neurophysiol. 2003;90: 415–430. doi: 10.1152/jn.01095.2002

3. Mann EO, Paulsen O. Role of GABAergic inhibition in hippocampal network oscillations. Trends Neurosci. 2007;30: 343–349. doi: 10.1016/j.tins.2007.05.003

4. Haider B. Neocortical Network Activity In Vivo Is Generated through a Dynamic Balance of Excitation and Inhibition. J Neurosci. 2006;26: 4535–4545. doi: 10.1523/JNEUROSCI.5297-05.2006

5. Isaacson JS, Scanziani M. How inhibition shapes cortical activity. Neuron. Elsevier Inc.; 2011;72: 231–243. doi:10.1016/j.neuron.2011.09.027

6. Buzsáki G, Watson BO. Brain rhythms and neural syntax: Implications for efficient coding of cognitive content and neuropsychiatric disease. Dialogues Clin Neurosci. 2012;14: 345–367. doi:10.1097/ALN.0b013e318212ba87

7. Buzsáki G. Rhythms of the Brain. Oxford: Oxford University Press; 2006. doi:10.1093/acprof:oso/9780195301069.001.0001

8. Buzsáki G. Theta oscillations in the hippocampus. Neuron. 2002;33: 325–340. doi:10.1016/S0896-6273(02)00586-X

9. Goutagny R, Jackson J, Williams S. Self-generated theta oscillations in the hippocampus. Nat Neurosci. 2009;12: 1491–3. doi:10.1038/nn.2440

10. Tort ABL, Scheffer-Teixeira R, Souza BC, Draguhn A, Brankák J. Theta-associated high-frequency oscillations (110-160Hz) in the hippocampus and neocortex. Prog Neurobiol. 2013;100: 1–14. doi:10.1016/j.pneurobio.2012.09.002

11. Pavlov I, Savtchenko LP, Song I, Koo J, Pimashkin A, Rusakov D a, et al. Tonic GABAA conductance bidirectionally controls interneuron firing pattern and synchronization in the CA3 hippocampal network. Proc Natl Acad Sci U S A. 2014;111: 504–9. doi:10.1073/pnas.1308388110

12. Mok SY, Nadasdy Z, Lim YM, Goh SY. Ultra-slow oscillations in cortical networks in vitro. Neuroscience. 2012;206: 17–24. doi: 10.1016/j.neuroscience.2012.01.009

13. Keren H, Marom S. Long-range synchrony and emergence of reentry in neural networks. Sci Rep. Nature Publishing Group; 2016;6: 1–17. doi: 10.1038/srep36837

14. Colombi I, Tinarelli F, Pasquale V, Tucci V, Chiappalone M. Corrigendum: A simplified in vitro experimental model encompasses the essential features of sleep [Front. Neurosci., 10, (2016), (315)] doi: 10.3389/fnins.2016.00315. Front Neurosci. 2016;10. doi:10.3389/fnins.2016.00409

15. Chiappalone M, Koudelka-hep M. Network dynamics and synchronous activity in cultured cortical neurons. Int J Neural Syst. 2007;17: 87–103. doi: 10.1142/S0129065707000968

16. Krahe R, Gabbiani F. Burst firing in sensory systems. Nat Rev Neurosci. 2004;5: 13–23. doi:10.1038/nrn1296

17. Huerta PT, Lisman JE. Bidirectional synaptic plasticity induced by a single burst during cholinergic theta oscillation in CA1 in vitro. Neuron. 1995;15: 1053–63. doi:http://dx.doi.org/10.1016/0896-6273(95)90094-2

18. Kim JH, Heo R, Choi JH, Lee KJ. Dynamic transitions among multiple oscillators of synchronized bursts in cultured neural networks. J Stat Mech Theory Exp. 2014;4: P04019. doi:10.1088/1742-5468/2014/04/P04019

19. Khazipov R, Sirota A, Leinekugel X, Holmes GL, Ben-Ari Y, Buzsáki G. Early motor activity drives spindle bursts in the developing somatosensory cortex. Nature. 2004;432: 758–761. doi:10.1038/nature03132

20. Leinekugel X, Khazipov R, Cannon R, Hirase H, Ben-Ari Y, Buzsáki G. Correlated bursts of activity in the neonatal hippocampus in vivo. Science. 2002;296: 2049–2052. doi:10.1126/science.1071111

21. Madhavan R, Chao ZC, Potter SM. Plasticity of recurring spatiotemporal activity patterns in cortical networks. Phys Biol. 2008;4: 181–193. doi: 10.1088/1478-3975/4/3/005.Plasticity

22. Wagenaar DA, Nadasdy Z, Potter SM. Persistent dynamic attractors in activity patterns of cultured neuronal networks. Phys Rev E - Stat Nonlinear, Soft Matter Phys. 2006;73: 1–17. doi:10.1103/PhysRevE.73.051907

23. Leondopulos SS, Boehler MD, Wheeler BC, Brewer GJ. Chronic stimulation of cultured neuronal networks boosts low frequency oscillatory activity at theta and gamma with spikes phase-locked to gamma frequencies. j Neural Eng. 2012;9: 1–20. doi:10.1088/1741-2560/9/2/026015.Chronic

24. Maeda E, Robinson HP, Kawana A. The mechanisms of generation and propagation of synchronized bursting in developing networks of cortical neurons. J Neurosci. 1995;15: 6834–6845. Available: http://www.jneurosci.org/content/15/10/6834

25. Maeda E, Kuroda Y, Robinson HP, Kawana a. Modification of parallel activity elicited by propagating bursts in developing networks of rat cortical neurones. Eur J Neurosci. 1998;10: 488–96. Available: http://www.ncbi.nlm.nih.gov/pubmed/9749711

26. Wagenaar D a, Pine J, Potter SM. An extremely rich repertoire of bursting patterns during the development of cortical cultures. BMC Neurosci. 2006;7: 11. doi: 10.1186/1471-2202-7-11

27. McCormick D, Contreras D. On the cellular and network bases of epileptic seizures. Annu Rev Physiol. 2001;63: 815–846. doi: 10.1146/annurev.physiol.63.1.815

28. Chiu C, Weliky M. Spontaneous activity in developing ferret visual cortex in vivo. J Neurosci. 2001;21: 8906–8914. Available: http://www.jneurosci.org/content/21/22/8906.long

29. Weliky M, Katz LC. Correlational structure of spontaneous neuronal activity in the developing lateral geniculate nucleus in vivo. Science. 1999;285: 599–604. doi: 10.1126/science.285.5427.599

30. Shahaf G, Eytan D, Gal A, Kermany E, Lyakhov V, Zrenner C, et al. Order-based representation in random networks of cortical neurons. PLoS Comput Biol. 2008;4: e1000228. doi:10.1371/journal.pcbi.1000228

31. Pimashkin A, Kastalskiy I, Simonov A, Koryagina E, Mukhina I, Kazantsev V. Spiking signatures of spontaneous activity bursts in hippocampal cultures. Front Comput Neurosci. 2011;5: 46. doi:10.3389/fncom.2011.00046

32. Pimashkin A, Gladkov A, Agrba E, Mukhina I, Kazantsev V. Selectivity of stimulus induced responses in cultured hippocampal networks on microelectrode arrays. Cogn Neurodyn. Springer Netherlands; 2016; 1–13. doi:10.1007/s11571-016-9380-6

33. Gandolfo M, Maccione a, Tedesco M, Martinoia S, Berdondini L. Tracking burst patterns in hippocampal cultures with high-density CMOS-MEAs. J Neural Eng. 2010;7: 056001. doi:10.1088/1741-2560/7/5/056001

34. Raichman N, Ben-Jacob E. Identifying repeating motifs in the activation of synchronized bursts in cultured neuronal networks. J Neurosci Methods. 2008;170: 96–110. doi:10.1016/j.jneumeth.2007.12.020

35. Idelson MS, Ben-Jacob E, Hanein Y. Innate Synchronous Oscillations in Freely-Organized Small Neuronal Circuits. PLoS One. 2010;5: 1–9. doi: 10.1371/journal.pone.0014443

36. Ito D, Tamate H, Nagayama M, Uchida T, Kudoh SN, Gohara K. Minimum neuron density for synchronized bursts in a rat cortical culture on multielectrode arrays. Neuroscience. Elsevier Inc.; 2010;171: 50–61. doi: 10.1016/j.neuroscience.2010.08.038

37. Ide N, Andruska A, Boehler M, Wheeler BC, Brewer GJ. Chronic network stimulation enhances evoked action potentials. J Neural Eng. 2010;7: 1–15. doi: 10.1088/1741-2560/7/1/016008

38. Koryagina EA, Pimashkin AS, Kazantsev VB, Mukhina IV. Dynamics of stimulated bioelectrical activity in neural networks in vitro. Vestn Lobachevsky Univ Nizhni Novgorod. 2011;2: 254–261.

39. Shirokova OM, Frumkina LE, Vedunova MV, Mitroshina EV, Zakharov YN, Khaspekov LG, et al. Morphofunctional Patterns of Neuronal Network Developing in Dissociated Hippocampal Cell Cultures. Sovrem Tehnol v Med. 2013;5: 6–12.

40. Chiappalone M, Bove M, Vato A, Tedesco M, Martinoia S. Dissociated cortical networks show spontaneously correlated activity patterns during in vitro development. Brain Res. 2006;1093: 41–53. doi:10.1016/j.brainres.2006.03.049

41. Van Pelt J, Corner M a., Wolters PS, Rutten WLC, Ramakers GJ a. Longterm stability and developmental changes in spontaneous network burst firing patterns in dissociated rat cerebral cortex cell cultures on multielectrode arrays. Neurosci Lett. 2004;361: 86–89. doi:10.1016/j.neulet.2003.12.062

42. Tokuda M, Kiyohara A, Taguch T, Kudoh SN. The effects of the current stimulation on electrical activity in dissociated neurons. 20th Anniversary MHS 2009 and Micro-Nano Global COE - 2009 International Symposium on Micro-NanoMechatronics and Human Science. 2009. pp. 118–122. doi: 10.1109/MHS.2009.5352068

43. Zhu G, Li X, Pu J, Chen W, Luo Q. Transient alterations in slow oscillations of hippocampal networks by low-frequency stimulations on multielectrode arrays. Biomed Microdevices. 2010;12: 153–158. doi:10.1007/s10544-009-9370-0

44. Gladkov AA, Pimashkin AS, Lepina AP, Kazantsev VB, Mukhina IV. Features of neural network response caused by electrical stimulation in mature hippocampal cell culture of mice. Vestn Lobachevsky state Univ Nizhni Novgorod. 2014;1: 57–64. Available: http://www.vestnik.unn.ru/en/nomera?anum_eng=8248

45. Frega M, Tedesco M, Massobrio P, Pesce M, Martinoia S. Network dynamics of 3D engineered neuronal cultures: a new experimental model for in-vitro electrophysiology. Sci Rep. 2014;4: 5489. doi:10.1038/srep05489

46. Gritsun T a., le Feber J, Rutten WLC. Growth Dynamics Explain the Development of Spatiotemporal Burst Activity of Young Cultured Neuronal Networks in Detail. PLoS One. 2012;7. doi:10.1371/journal.pone.0043352

47. Chen X, Dzakpasu R. Observed network dynamics from altering the balance between excitatory and inhibitory neurons in cultured networks. Phys Rev E - Stat Nonlinear, Soft Matter Phys. 2010;82: 1–8. doi: 10.1103/PhysRevE.82.031907

48. Huang YT, Cheung YL, Song H, Lai PY, Chan CK. 8th Int. Meeting on Substrate-Integrated Microelectrode Arrays. Spontaneous reverberation in developing neuronal culture networks. Reutlingen, Germany; 2012. pp. 86–87.

49. Robinson E. Hippocampal rhythmic slow activity (RSA, theta): A critical analysis of selected studies and discussion of possible species-differences. Brain Res. 1980;2: 69–101. doi: http://dx.doi.org/10.1016/0165-0173(80)90004-1

50. Quiroga RQ, Nadasdy Z, Ben-Shaul Y. Unsupervised spike detection and sorting with wavelets and superparamagnetic clustering. Neural Comput. 2004;16: 1661–87. doi:10.1162/089976604774201631

51. Vedunova M, Sakharnova T, Mitroshina E, Perminova M, Pimashkin A, Zakharov Y, et al. Seizure-like activity in hyaluronidase-treated dissociated hippocampal cultures. Front Cell Neurosci. 2013;7: 149. doi: 10.3389/fncel.2013.00149

52. Davies DL, Bouldin DW. A cluster separation measure. IEEE Trans Pattern Anal Mach Intell. 1979;1: 224–227. doi:10.1109/TPAMI.1979.4766909

53. Penttonen M, Buzsáki G. Natural logarithmic relationship between brain oscillators. Thalamus Relat Syst. 2003;2: 145–152. doi:10.1016/S1472-9288(03)00007-4

54. Sirota A, Csicsvari J, Buhl D, Buzsáki G. Communication between neocortex and hippocampus during sleep in rodents. Proc Natl Acad Sci U S A. 2003;100: 2065–2069. doi:10.1073/pnas.0437938100

55. Kopell N, Ermentrout GB, Whittington M a, Traub RD. Gamma rhythms and beta rhythms have different synchronization properties. Proc Natl Acad Sci U S A. 2000;97: 1867–1872. doi:10.1073/pnas.97.4.1867

56. Sullivan D, Csicsvari J, Mizuseki K, Montgomery S, Diba K, Buzsáki G. Relationships between hippocampal sharp waves, ripples, and fast gamma oscillation: influence of dentate and entorhinal cortical activity. J Neurosci. 2011;31: 8605–8616. doi:10.1523/JNEUROSCI.0294-11.2011

57. Jackson J, Goutagny R, Williams S. Fast and slow gamma rhythms are intrinsically and independently generated in the subiculum. J Neurosci. 2011;31: 12104–12117. doi: 10.1523/JNEUROSCI.1370-11.2011

58. Soltesz I, Deschenes M. Low- and high-frequency membrane potential oscillations during theta activity in CA1 and CA3 pyramidal neurons of the rat hippocampus under ketamine-xylazine anesthesia. J Neurophysiol. 1993;70: 97–116. Available: http://jn.physiology.org/content/70/1/97

59. Richards KL, Kurniawan ND, Kim TH, Keller MD, Ullmann JFP, Cole S, et al. Hippocampal volume and cell density changes in a mouse model of human genetic epilepsy. Neurology. 2013;80: 1240–1246. doi: 10.1212/WNL.0b013e31828970ec

60. Kiyohara A, Taguchi T, Kudoh SN. Effects of Electrical Stimulation on Autonomous Electrical Activity in a Cultured Rat Hippocampal Neuronal Network. EEJ Trans Electr Electron Eng. 2011;18: 163–167. doi: 10.1002/tee.20639

61. Bao W, Wu J-Y. Propagating wave and irregular dynamics: spatiotemporal patterns of cholinergic theta oscillations in neocortex in vitro. J Neurophysiol. 2003;90: 333–341. doi:10.1152/jn.00715.2002

62. Niedringhaus M, Chen X, Dzakpasu R, Conant K. MMPs and Soluble ICAM-5 Increase Neuronal Excitability within In Vitro Networks of Hippocampal Neurons. PLoS One. 2012;7: 1–9. doi: 10.1371/journal.pone.0042631

63. Ito D, Komatsub T, Goharab K. Measurement of saturation processes in glutamatergic and GABAergic synapse densities during long-term development of cultured rat cortical networks. Brain Res. 2013;1534: 22–32. Available: http://doi.org/10.1016/j.brainres.2013.08.004

64. Benson DL, Cohen P a. Activity-independent segregation of excitatory and inhibitory synaptic terminals in cultured hippocampal neurons. J Neurosci. 1996;16: 6424–32. Available: http://www.ncbi.nlm.nih.gov/pubmed/8815921

65. Benson DL, Tanaka H. N-Cadherin Redistribution during Synaptogenesis in Hippocampal Neurons. J Neurosci. 1998;18: 6892–6904. Available: http://www.jneurosci.org/content/18/17/6892.long

66. Yamada MK, Nakanishi K, Ohba S, Nakamura T, Ikegaya Y, Nishiyama N, et al. Brain-derived neurotrophic factor promotes the maturation of GABAergic mechanisms in cultured hippocampal neurons. J Neurosci. 2002;22: 7580–5. doi:22/17/7580 [pii]

67. Reimer T, Baumann W, Gimsa J. Population Bursts of Parvalbumin-Positive Interneurons Inhibit Spiking Pyramidal Cells in Spontaneously Active Cortical. 2012;6: 1033–1042. Available: http://www.davidpublishing.com/davidpublishing/Upfile/1/11/2013/2013011102854784.pdf

68. Stephens CL, Toda H, Palmer TD, Demarse TB, Ormerod BK. Adult neural progenitor cells reactivate superbursting in mature neural networks. Exp Neurol. Elsevier Inc.; 2012;234: 20–30. doi:10.1016/j.expneurol.2011.12.009

69. Wagenaar DA. Development and Control of Epileptiform Bursting in Dissociated Cortical Cultures. 2006. Dissertation (Ph.D.), California Institute of Technology. http://resolver.caltech.edu/CaltechETD:etd-07032005-170918

70. Konopacki J, Kowalczyk T, Golebiewski H, Eckersdorf B, Konopacki J. Window effect of temperature on carbachol-induced theta-like activity recorded in hippocampal formation in vitro. Brain Res. 2001;901: 184–194. doi: 10.1016/S0006-8993(01)02355-1

71. Konopacki J, Gdqbiewski H, Eckersdorf B. In vitro recorded theta-like activity in the limbic cortex: Comparison with spontaneous theta and epileptiform discharges. Acta Neurobiol Exp (Wars). 2000;60: 67–85. Available: http://www.ane.pl/linkout.php?vol=60&no=1&fpp=67

72. Gritsun TA, Le Feber J, Stegenga J, Rutten WLC. Network bursts in cortical cultures are best simulated using pacemaker neurons and adaptive synapses. Biol Cybern. Springer Berlin / Heidelberg; 2010;102: 293–310. doi: 10.1007/s00422-010-0366-x

